# RBPMetaDB: A comprehensive annotation of mouse RNA-Seq datasets with perturbations of RNA-binding proteins

**DOI:** 10.1101/326116

**Authors:** Jin Li, Su-Ping Deng, Jacob Vieira, James Thomas, Valerio Costa, Ching-San Tseng, Franjo Ivankovic, Alfredo Ciccodicola, Peng Yu

**Affiliations:** Department of Electrical and Computer Engineering, College Station, TX 77843, USA.; TEES-AgriLife Center for Bioinformatics and Genomic Systems Engineering, Texas A&M University, College Station, TX 77843, USA.; The Department of Microbiology, University of Massachusetts Amherst, Amherst, MA, USA.; Department of Molecular Genetics and Microbiology, Center for NeuroGenetics and the Genetics Institute, College of Medicine, University of Florida, Gainesville, Florida, USA.; Institute of Genetics and Biophysics “Adriano Buzzati-Traverso”, Consiglio Nazionale delle Ricerche, Via P. Castellino 111, 80131 Naples, Italy.; Institute of Cellular and Organismic Biology, Academia Sinica, Taipei, 11529, Taiwan.; Department of Science and Technology, University Parthenope of Naples, 80131 Naples, Italy.

## Abstract

RNA-binding proteins may play a critical role in gene regulation in various diseases or biological processes by controlling post-transcriptional events such as polyadenylation, splicing, and mRNA stabilization via binding activities to RNA molecules. Due to the importance of RNA-binding proteins in gene regulation, a great number of studies have been conducted, resulting in a large amount of RNA-Seq datasets. However, these datasets usually do not have structured organization of metadata, which limits their potentially wide use. To bridge this gap, the metadata of a comprehensive set of publicly available mouse RNA-Seq datasets with perturbed RNA-binding proteins were collected and integrated into a database called RBPMetaDB. This database contains 278 mouse RNA-Seq datasets for a comprehensive list of 163 RNA-binding proteins. These RNA-binding proteins account for only ∼10% of all known RNA-binding proteins annotated in Gene Ontology, indicating that most are still unexplored using high-throughput sequencing. This negative information provides a great pool of candidate RNA-binding proteins for biologists to conduct future experimental studies. In addition, we found that DNA-binding activities are significantly enriched among RNA-binding proteins in RBPMetaDB, suggesting that prior studies of these DNA- and RNA-binding factors focus more on DNA-binding activities instead of RNA-binding activities. This result reveals the opportunity to efficiently reuse these data for investigation of the roles of their RNA-binding activities. A web application has also been implemented to enable easy access and wide use of RBPMetaDB. It is expected that RBPMetaDB will be a great resource for improving understanding of the biological roles of RNA-binding proteins.

Database URL: http://rbpmetadb.yubiolab.org

## Introduction

A lack of fully structured metadata limits the wide use of valuable RNA-Seq datasets in public repositories such as Gene Expression Omnibus (GEO) (1) and ArrayExpress (2). To fill this gap, manual curation has been shown to be an effective way to collect data resources (3) and has been applied to develop and maintain metadata databases(4). For example, microarray and RNA-Seq datasets have been curated for the downstream analyses in Expression Atlas (5) and in epidermal development(6). We previously launched two databases, RNASeqMetaDB (7) and SFMetaDB (8), to facilitate access to the metadata of publicly available mouse RNA-Seq datasets with perturbed disease-related genes and splicing factors, respectively. Here, we present a new database, RBPMetaDB, for the metadata of RNA-Seq datasets with perturbed RNA-binding proteins (RBPs).

RBPs play a critical role in multiple cellular processes in eukaryotes. RBPs bind to double- or single-stranded RNA molecules and are potential key factors in biological processes, such as pre-mRNA splicing, RNA methylation, and protein translation (9). Besides influencing each of these processes, RBPs also provide a link between them (10). The perturbation of these intricate networks can destroy the coordination of complex post-transcriptional events and lead to disease (11).

According to recent genomic data and evidence derived from animal models, RBPs play a crucial role in the pathogenesis of many complex human diseases, including neurological disorders (12), Mendelian diseases(13), and cancer (14). These diseases have been demonstrated to have strong associations with aberrant functions or expression of RBPs, which can impact many different genes and pathways. Some diseases can be caused by loss of function of RBPs, such as Fragile X syndrome, paraneoplastic neurologic syndromes, and spinal muscular atrophy (9). For example, Fragile X syndrome can be caused by the deficiency of gene fragile X mental retardation (*FMR1*)(15). Alternatively, some diseases can be caused by gain of function of RBPs, including myotonic dystrophy, Fragile X tremor ataxia syndrome, and oculopharyngeal muscular dystrophy (OPMD) (9). For instance, OPMD is generated by the accumulation of aggregates in the nuclei of skeletal muscle fibers caused by mutants in the protein PABPN1 (16). And a deficiency of PABPN1 can induce progressive muscle weakness in muscular dystrophy(17).

To investigate the functions of RBPs in biological processes or diseases such as the ones mentioned above, a large number of studies have been conducted, resulting in exponential growth of RBP-related papers in recent years (Figure 1). For example, more than 1,000 papers were published in 2017 alone. Among the studies on RBPs, a large number of RNA-Seq datasets have been generated in loss- or gain-of-function experiments and are publicly available from online repositories like GEO (1). However, because GEO does not have a stringent requirement for metadata of the submitted datasets, the metadata are non-uniformly maintained across different datasets, resulting in inconsistent dataset annotation and sometimes ambiguity. Such a deficiency makes it difficult to identify useful datasets with high precision and recall, which limits the wide use of the datasets.

**Figure 1.**
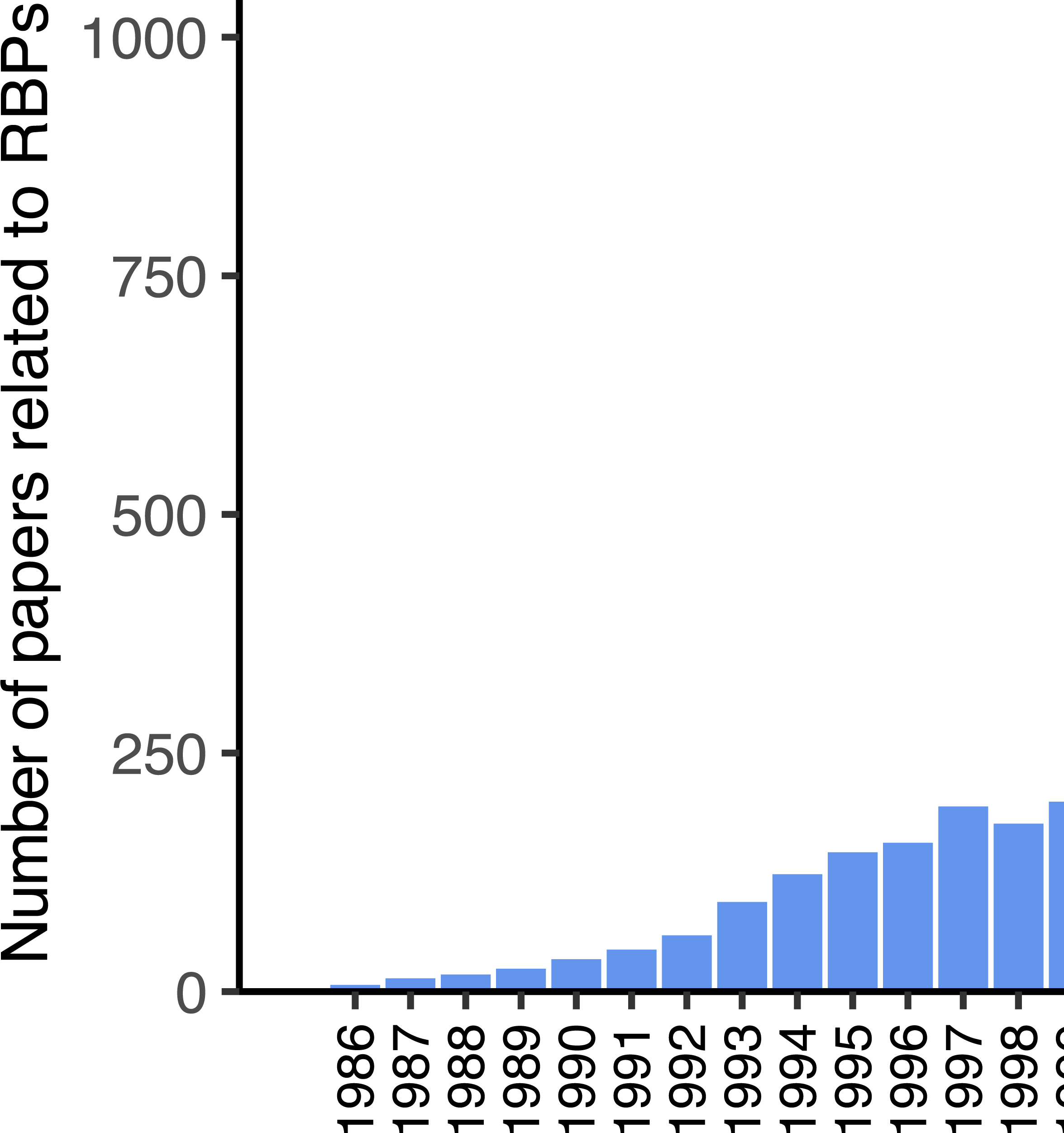
The rapid growth of papers related to RPBs in PubMed. Approximately 10,000 papers related to RPBs are indexed on PubMed according to the query of “RNA binding protein”[tiab] OR “RNA binding proteins” [tiab] at the time of writing. Since 2012, the number of papers published per year has been increasing more rapidly than ever before. In 2017 alone, over 1,000 papers were published.

To address this challenge, we worked to curate RNA-Seq datasets from GEO and ArrayExpress with one or more RBPs being perturbed, e.g., by knock-out, knock-down, or overexpression. Important dataset annotations such as genotypes and PubMed references were manually curated to ensure high accuracy. Curated datasets can be used in gene expression analysis (18,19) and alternative splicing analysis (20,21) for biological hypothesis generation (11) via a signature comparison approach (22). To facilitate the use of our curated datasets, the metadata information of these datasets was imported into a database called RBPMetaDB. It should be mentioned that our database differs greatly from Expression Atlas in the sense that the latter is not mainly about datasets where specific genes are perturbed and so are not guaranteed to be complete in this aspect.

In this paper, we describe our main curation methods used in constructing RBPMetaDB and the statistics of the database. To demonstrate the use of RBPMetaDB, a number of promising candidate RBPs have been identified by comparing RBPs with RNA-Seq datasets and all the RBPs annotated in Gene Ontology (GO). In addition, a web application has been developed to host the database to broaden the use of curated metadata and the original raw datasets among biomedical communities.

## Materials and methods

### Metadata curation of GEO/ArrayExpress RNA-Seq datasets and RBPMetaDB web application deployment

To collect RNA-Seq datasets for RBPs from GEO comprehensively, we first extracted 1,587 mouse RBPs annotated in GO (accession GO:0003723) (23). Each of these RBPs was queried against GEO for mouse RNA-Seq data using the query `(<official_symbol>[Title] OR <official_symbol>[Description]) NOT SuperSeries[Description] AND gse[Entry Type] AND` “`Mus musculus`” `[porgn:__txid10090] AND (`“`expression profiling by high throughput sequencing`” `[DataSet Type] OR`” `non coding rna profiling by high throughput sequencing″[DataSet Type])` and against ArrayExpress using the query `(<official_symbol> AND organism:` “ `Mus musculus`” `AND exptype:`” `sequencing assay`” `AND exptype:` “ `rna assay`” `).”`). These queries resulted in 1,194 unique datasets in mice. Due to the limitations of the search functions of GEO and ArrayExpress, many of these datasets do not have perturbed RBPs despite the official symbols of some RBPs being mentioned in the titles or descriptions of the datasets. To retain the datasets with perturbations of RBPs, we manually curated each dataset (24) and retained datasets with biological replications per comparison condition, with at least one RBP being knocked-out, knocked-down, or overexpressed (along with the corresponding wild-type or control samples) in mice. For the datasets that do not have associated PubMed IDs on GEO and ArrayExpress, we manually added the PubMed IDs.

To facilitate access to these datasets, we launched a database called RBPMetaDB (http://rbpmetadb.yubiolab.org). RBPMetaDB is implemented using Flask (http://flask.pocoo.org), a microframework for web development in Python. The MySQL database is used for data storage. The website of RBPMetaDB is freely available, and it presents the GEO/ArrayExpress accession numbers, descriptions, number of samples, associated curated RBPs, perturbation, and PubMed references for each RNA-Seq dataset.

### Domain structure analysis of RBPs

Protein domain structure analysis of RBPs was performed to identify critical RBPs for future studies. First, all RBPs annotated to the “RNA binding” GO term (GO: 0003723) were retrieved using the R package GO.db (25). Using the UniProt annotation of the Pfam families assigned to the RBP protein domains (26), the number of RBPs with specific Pfam families was calculated using RBPs with the curated RNA-Seq datasets and using the total RBPs, respectively. To investigate the RNA binding effect, the number of RBPs of Pfam families with RNA binding activity was calculated, where RNA-related Pfam families were searched using the RESTful interface in the Pfam database. Figure 3 plots the number of RBPs with Pfam families specific to RNA binding for the RBPs with RNA-Seq data and all the RBPs. By comparing the domain families of the RBPs with RNA-Seq datasets to those of all the RBPs, the RBPs in relatively less-studied domain families can be promising candidates for future RBP studies.

## Results

### Data statistics

RBPMetaDB has 292 RNA-Seq datasets with 187 perturbed RBPs (**Table S1**), which account for only ∼10% of all annotated RBPs. Among these 187 RBPs, over 30% of them have more than one corresponding RNA-Seq dataset. Approximately 90% of datasets in RBPMetaDB have only one perturbed RBP, meaning that most studies are small-scale and well-focused. Also, RBPs with RNA-Seq data tend to have DNA-binding activity. To systematically examine the DNA-binding activity of RBPs, the GO term “DNA binding” (GO:0003677) was used to extract the genes with DNA-binding activity. By overlapping with DNA-binding proteins, 66 RBPs with RNA-Seq datasets and 207 RBPs without RNA-Seq datasets were shown to have DNA-binding activity. Taking the total 1,587 RBPs as background, Fisher’s exact test showed an enrichment of DNA-binding activity in RBPs with datasets compared to RBPs without datasets (*p*-value< 5.8 × 10^−14^). Specifically, for RBPs in RBPMetaDB, the proportion between RBPs with and without DNA-binding activity is 0.68 (66 over 97). On the contrary, the proportion of RBPs that do not have RNA-Seq datasets is only 0.17 (207 over 1,219). This large difference suggests that many datasets in RBPMetaDB were collected for their DNA-binding activity instead of RNA-binding activity, and these datasets are likely to be underanalyzed for RNA-binding activity, providing a cost-effective opportunity to reanalyze these datasets to study their related RNA biology. For example, *Ezh2* is the most-studied gene, with 35 RNA-Seq datasets in RBPMetaDB. However, most studies of EZH2, as a catalytic subunit of Polycomb Repressive Complex 2 (PRC2), focus on its capacity for mono-, di-, and trimethylation of histone H3 on lysine K27 (H3K27me1/2/3) (27).

Figure 2a shows that the main RBP perturbation type of all the datasets in RBPMetaDB, is knock-out (∼67%). The rest is knock-down (∼18%), overexpression (∼9%), knock-in (∼3.5%), and other (∼2.8%, e.g., treated with inhibitors or point mutation). Figure 2b shows that the US and Europe dominate the generation of RNA-Seq datasets for studying RBPs, with contributions of 60.1% and 23.4% of all the datasets, respectively. In addition, Figure 2c shows an increasing number of papers published about the RNA-Seq datasets in RBPMetaDB from 2010 to 2017. This increasing research interest worldwide will stimulate more investigation on RBPs.

**Figure 2.**
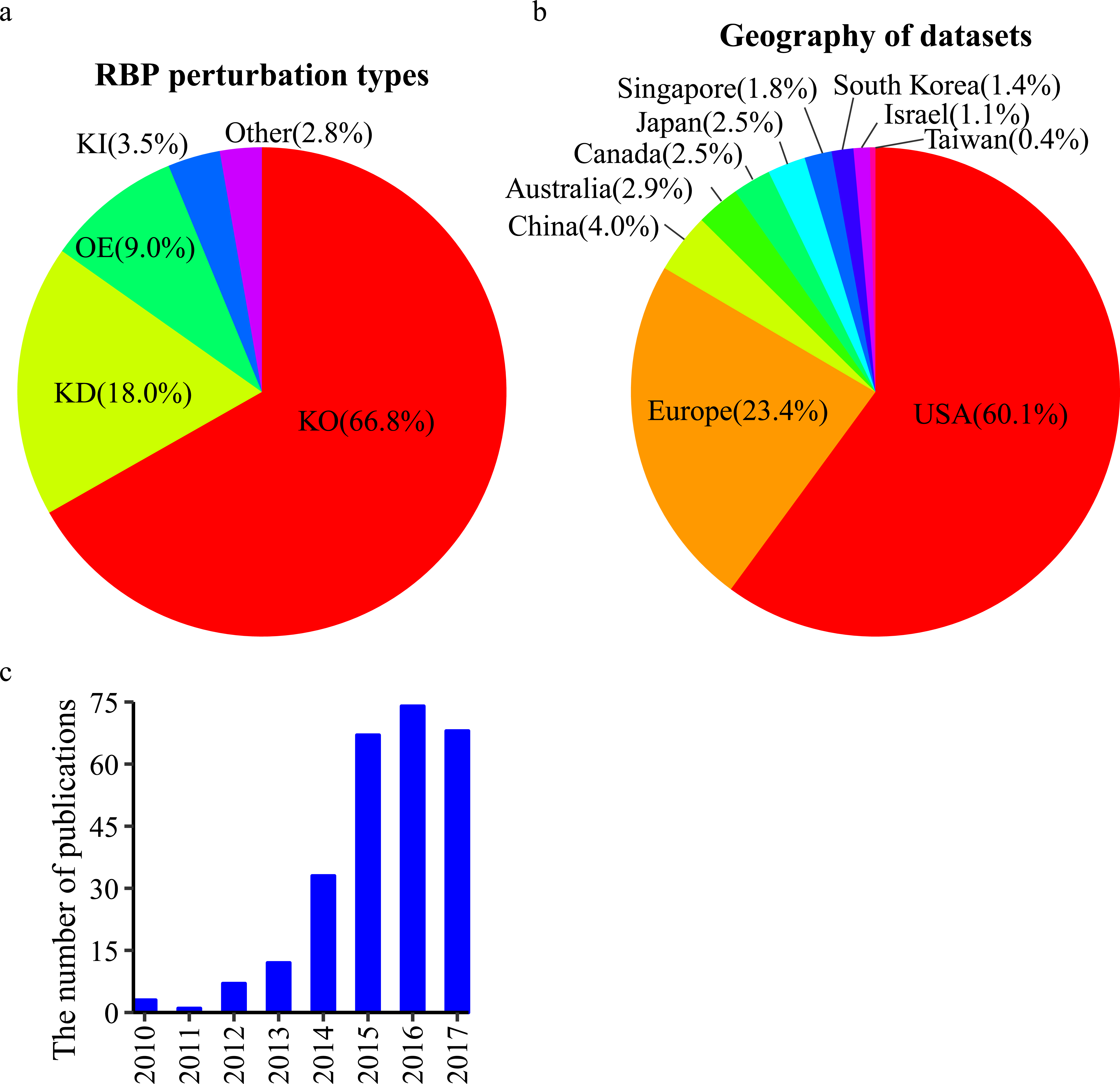
Statistics of curated RNA-Seq datasets for RBPs. (a) The distribution of perturbation types: knock-out (KO), knock-down (KD), overexpression (OE), knock-in (KI), and other (e.g., point mutations of RBPs or treatment with inhibitors of RBPs) among all the curated datasets. The percentages are shown between parentheses. Knock-out experiments are the most common. (b) The curated datasets are generated from research labs worldwide. The US is the dominant country with a contribution of 60.1% of all the datasets. (c) The number of associated publications for the datasets increased from 2010 to 2017. The slow-down of increase in 2016 and the drop in 2017 are likely due to the missing PMIDs annotation for a subset of the recently released datasets on GEO.

**Figure 3.**
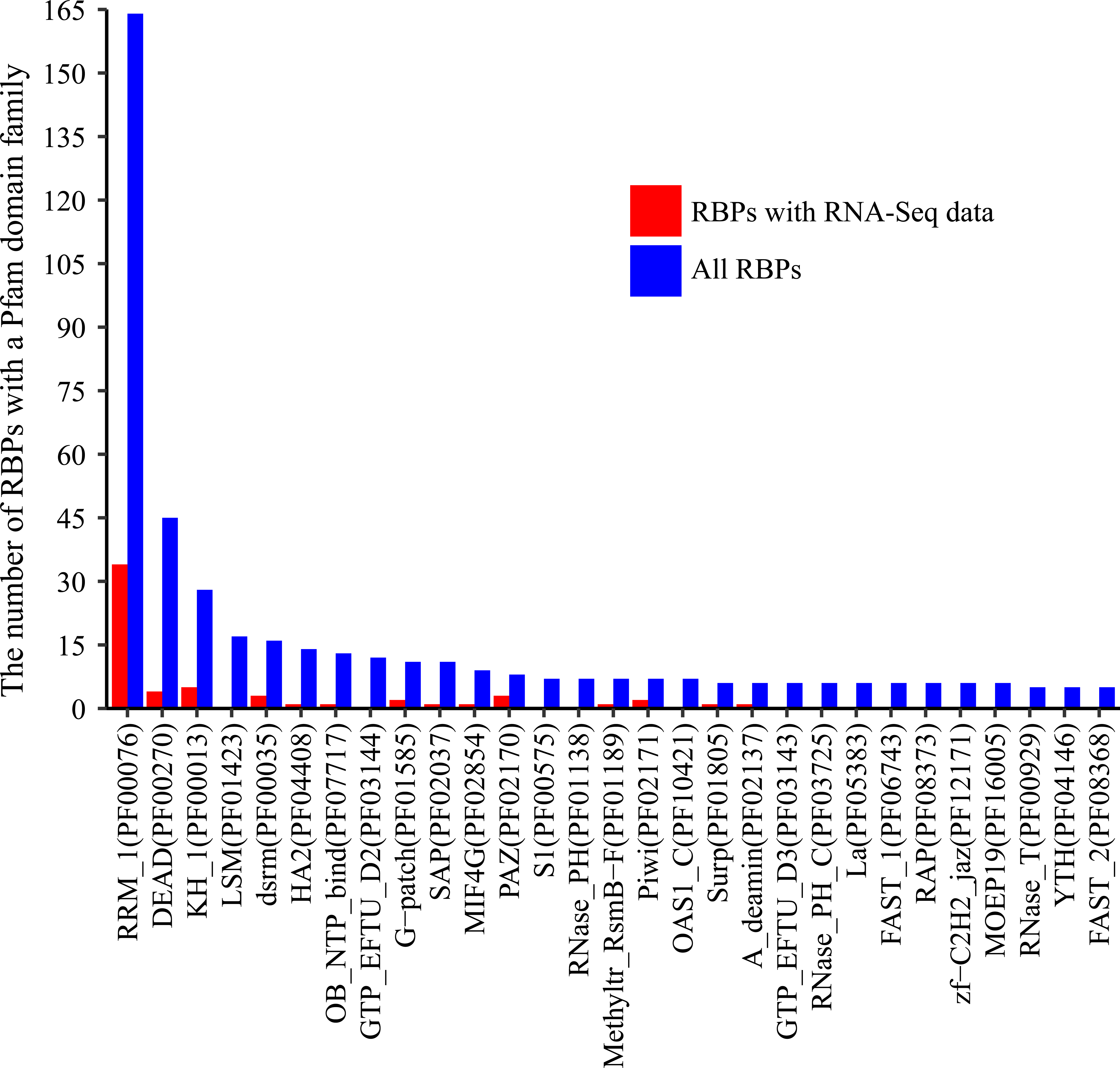
The number of RBPs containing a domain from a Pfam family with RNA-binding activity. Blue bars indicate the number of RBPs containing a domain from a family among all RBPs, and red bars indicate the numbers of RBPs containing a domain from a family among the RBPs with associated RNA-Seq datasets. Only families with a blue bar with > 5 RBPs are shown.

### Comparison of RBPs using protein domain analysis

Protein domains, as conserved protein structural units, typically characterize certain functional aspects of a protein, and proteins sharing similar domains tend to share similar functions. Since RBPs bind to RNAs, they should have RNA-binding domain. We therefore extracted the domain family information of all the RBPs according to Pfam domain family annotation (Finn et al., 2016). Figure 3 shows the protein domain families ordered by the number of RBPs with a domain from a Pfam family, and only families with RNA-binding activities with ≥ 5 annotated RBPs are shown. The most dominant domain family is RRM_1 (RNA recognition motif, PF00076) and the RBPs with domains from this family are relatively well-studied (34 in RBPMetaDB over all 164 annotated). RBPs with domains from four additional families are fairly well-studied, including DEAD (DEAD/DEAH box helicase, PF00270), KH_1 (KH domain, PF00013), dsrm (double-stranded RNA-binding motif, PF00035), and HA2 (helicase-associated domain, PF04408). However, none of the RBPs with domains from two highly dominant domain families, LSM (PF01423) and GTP_EFTU_D2 (PF03144), has related RNA-Seq datasets yet, and they may be good candidates for future high-throughput sequencing studies.

What’s more, among the RBPs without related RNA-Seq datasets, 140 RBPs already have one or more mouse models (**Table S1**) on the International Mouse Strain Resource (IMSR) (28). For example, the gene Cleavage Stimulation Factor Subunit 2 Tau Variant (*Cstf2t*) has been demonstrated to be an important stage-specific regulator of *Crem* mRNA processing that controls *Crem* polyadenylation in mouse testis. *Cstf2t* can lead to an overall decrease of the Crem mRNAs generated from internal promoters in *Cstf2t*^-/-^ mice (29,30). Therefore, these 140 RBPs can be promising candidates for RNA-Seq studies in the future.

### Web interface

To facilitate the use of RBPMetaDB, a user-friendly website has been launched. The website allows users to access all the key information related to the curated RNA-Seq datasets, including the GEO/ArrayExpress accession numbers, dataset titles, numbers of samples, associated RBPs, perturbation types, and PubMed IDs (Figure 4). The contents in these fields are linked to the corresponding entries in GEO/ArrayExpress, metadata information for each dataset, MGI gene symbol, and PubMed, respectively. In the table view of the website, the first 10 entries are shown by default, but the user can easily select the number of entries to be visualized from a pop-up menu on the left side (Label A). Each table has six columns about the metadata in RBPMetaDB (Label B), and all columns can be sorted in ascending or descending order by clicking column headers. The search boxes at the bottom of all the fields support field-specific search by regular expression (Label C). For example, to search for multiple gene symbols in the “RNA binding proteins” column, one can specify the gene symbols joined by “|”By searching a gene of interest, users can find all RNA-Seq datasets with the gene perturbed. Take as an example METTL3, which is an important enzyme involved in the post-transcriptional methylation of internal adenosine residues in eukaryotic mRNA (31)— it can be demonstrated that RPBMetaDB greatly outperforms GEO in terms of search efficiency. When the keyword “;Mettl3” is searched on RBPMetaDB, it returns six highly accurate mouse RNA-Seq datasets from *Mettl3* loss- or gain-of-function studies (Figure 5a). GEO returns 35 mouse RNA-Seq datasets with the query of “Mettl3” in dataset titles and descriptions (Figure 5b), but it is impossible to directly identify which RNA-Seq datasets are from loss- or gain-of-function experiments of *Mettl3*. On the contrary, RBPMetaDB does not return irrelevant datasets of a given RBP, and it returns more accurate results than GEO.

**Figure 4.**
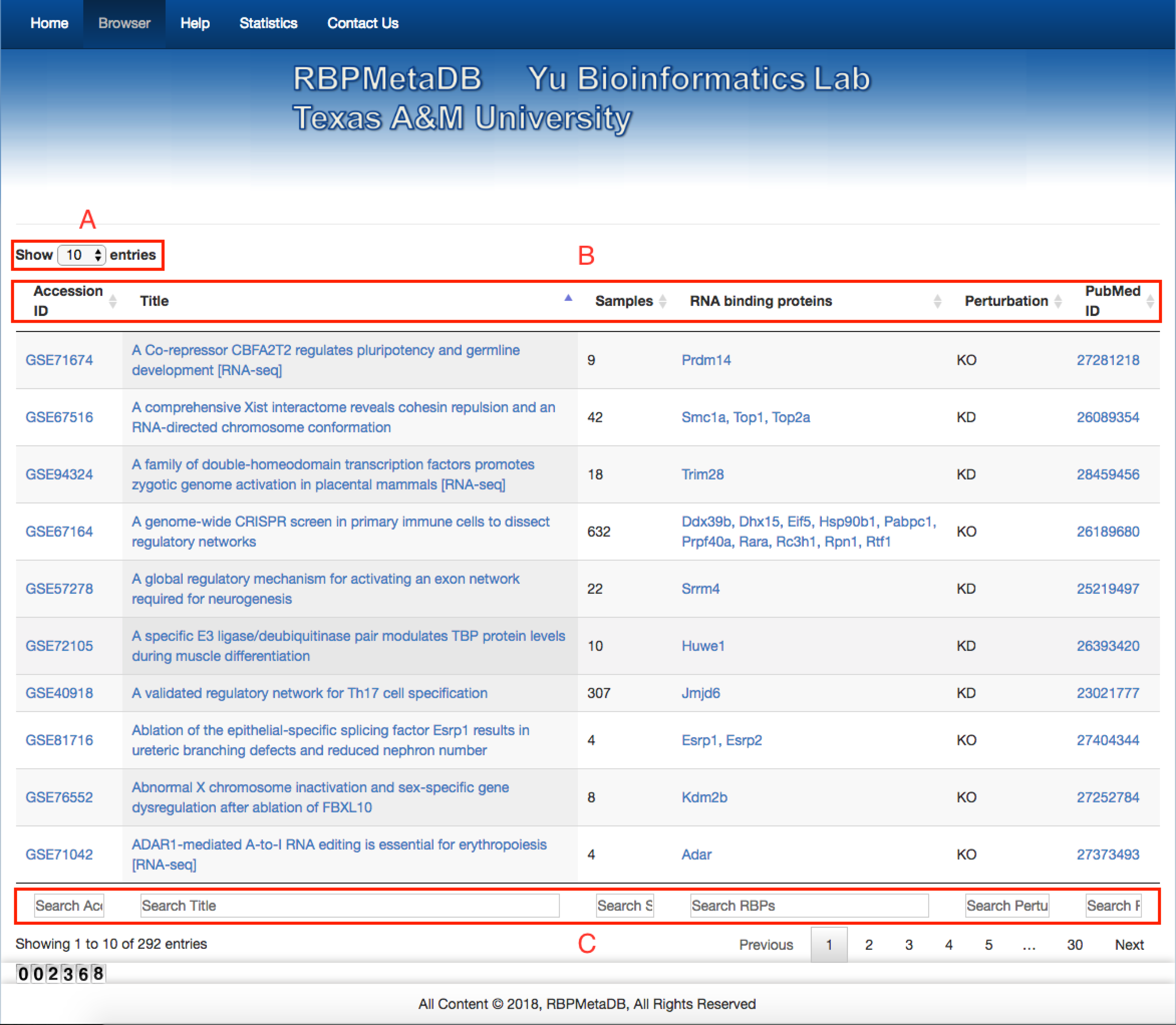
Web interface of RBPMetaDB. The RBPMetaDB website presents information about the mouse RNA-Seq datasets with perturbed RBPs. Label A refers to the maximum number of entries shown on a page. Label B is about the relevant information for each RNA-Seq dataset including GEO accession numbers, titles of the datasets in GEO, number of samples, official gene symbols from Mouse Genome Informatics (MGI), perturbation types of the RBPs associated with a dataset, and PMIDs of the related papers. Label C refers to the field specific search boxes.

**Figure 5.**
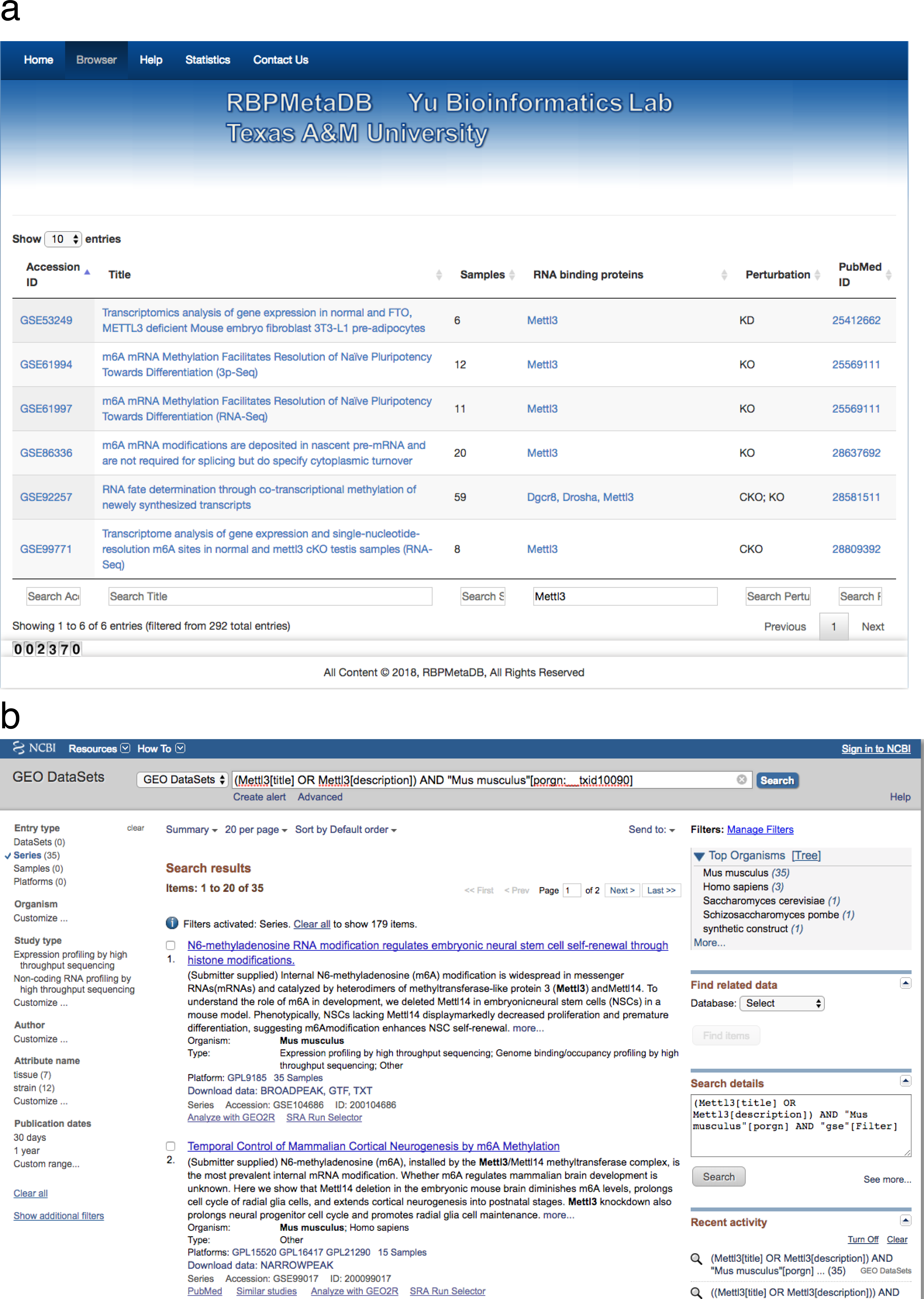
A use case of RBPMetaDB for the mouse RPB METTL3. (a) Here is a use case of RPB METTL3 to demonstrate the advantage of RBPMetaDB over GEO. By using the keyword “Mettl3,” RBPMetaDB accurately returns six mouse RNA-Seq datasets with *Mettl3* perturbed. (b) However, GEO returns 35 mouse RNA-Seq datasets without identifying which datasets are from experiments with *Mettl3* perturbed.

## Conclusions

RBPMetaDB provides the first comprehensive, manually curated database of mouse RNA-Seq datasets with specific RBPs being perturbed. At the time of writing, it consists of the annotation of 292 mouse RNA-Seq datasets. These datasets provide valuable information for studying RNA-binding activity of RBPs. To keep RBPMetaDB updated, every six months, we will extract an updated list of RBPs from GO annotation to search GEO and ArrayExpress for newly released mouse RNA-Seq datasets and curate them. The results will be added to RBPMetaDB. RBPMetaDB will provide a valuable resource for many different research communities to understand how RBPs are involved in a variety of biological or disease processes.

## Acknowledgements

The authors thank Antonio Federico for his contribution to RBPMetaDB.

## References

1. Barrett, T., Wilhite, S.E., Ledoux, P., et al. (2013) NCBI GEO: archive for functional genomics data sets-update. Nucleic Acids Research, 41, D991– D995.

2. Kolesnikov, N., Hastings, E., Keays, M., et al. (2015) ArrayExpress update-simplifying data submissions. Nucleic Acids Research, 43, D1113–D1116.

3. Li, J., Deng, S.-P., Wei, G., et al. (2018) CITGeneDB: a comprehensive database of human and mouse genes enhancing or suppressing cold-induced thermogenesis validated by perturbation experiments in mice. Database, 2018, bay012–bay012.

4. Qin, B., Zhou, M., Ge, Y., et al. (2012) CistromeMap: a knowledgebase and web server for ChIP-Seq and DNase-Seq studies in mouse and human. Bioinformatics, 28, 1411–1412.

5. Petryszak, R., Keays, M., Tang, Y.A., et al. (2016) Expression Atlas update- an integrated database of gene and protein expression in humans, animals and plants. Nucleic Acids Res, 44, D746–752.

6. Li, J., Zheng, L., Uchiyama, A., et al. (in press) A data mining paradigm for identifying key factors in biological processes using gene expression data. Sci Rep.

7. Guo, Z., Tzvetkova, B., Bassik, J.M., et al. (2015) RNASeqMetaDB: a database and web server for navigating metadata of publicly available mouse RNA-Seq datasets. Bioinformatics, 31, 4038–4040.

8. Li, J., Tseng, C.S., Federico, A., et al. (2017) SFMetaDB: a comprehensive annotation of mouse RNA splicing factor RNA-Seq datasets. Database (Oxford), 2017.

9. Lukong, K.E., Chang, K.W., Khandjian, E.W., et al. (2008) RNA-binding proteins in human genetic disease. Trends Genet, 24, 416–425.

10. Glisovic, T., Bachorik, J.L., Yong, J., et al. (2008) RNA-binding proteins and post-transcriptional gene regulation. FEBS Lett, 582, 1977–1986.

11. Li, J., Yu, P. (2018) Genome-wide transcriptome analysis identifies alternative splicing regulatory network and key splicing factors in mouse and human psoriasis. Sci Rep, 8, 4124.

12. Zhou, H., Mangelsdorf, M., Liu, J., et al. (2014) RNA-binding proteins in neurological diseases. Sci China Life Sci, 57, 432–444.

13. Castello, A., Fischer, B., Hentze, M.W., et al. (2013) RNA-binding proteins in Mendelian disease. Trends Genet, 29, 318–327.

14. Pereira, B., Billaud, M., Almeida, R. (2017) RNA-Binding Proteins in Cancer: Old Players and New Actors. Trends Cancer, 3, 506–528.

15. Mientjes, E.J., Nieuwenhuizen, I., Kirkpatrick, L., et al. (2006) The generation of a conditional Fmr1 knock out mouse model to study Fmrp function in vivo. Neurobiol Dis, 21, 549–555.

16. Abu-Baker, A., Rouleau, G.A. (2007) Oculopharyngeal muscular dystrophy: recent advances in the understanding of the molecular pathogenic mechanisms and treatment strategies. Biochimica et biophysica acta, 1772, 173–185.

17. Dion, P., Shanmugam, V., Gaspar, C., et al. (2005) Transgenic expression of an expanded (GCG)13 repeat PABPN1 leads to weakness and coordination defects in mice. Neurobiol Dis, 18, 528–536.

18. Cress, W.D., Yu, P., Wu, J. (2017) Expression and alternative splicing of the cyclin-dependent kinase inhibitor-3 gene in human cancer. The international journal of biochemistry & cell biology, 91, 98–101.

19. Osenberg, S., Karten, A., Sun, J., et al. (2018) Activity-dependent aberrations in gene expression and alternative splicing in a mouse model of Rett syndrome. Proc Natl Acad Sci U S A.

20. Yu, P., Shaw, C.A. (2014) An efficient algorithm for accurate computation of the Dirichlet-multinomial log-likelihood function. Bioinformatics, 30, 1547–1554.

21. Dai, L., Chen, K., Youngren, B., et al. (2016) Cytoplasmic Drosha activity generated by alternative splicing. Nucleic Acids Res, 44, 10454–10466.

22. Li, Z., Li, J., Yu, P. (2017) l1kdeconv: an R package for peak calling analysis with LINCS L1000 data. BMC Bioinformatics, 18, 356.

23. Ashburner, M., Ball, C.A., Blake, J.A., et al. (2000) Gene ontology: tool for the unification of biology. The Gene Ontology Consortium. Nat Genet, 25, 25–29.

24. Li, Z., Li, J., Yu, P. (2018) GEOMetaCuration: a web-based application for accurate manual curation of Gene Expression Omnibus metadata. Database, 2018, bay019–bay019.

25. Carlson, M. (2017) GO.db: A set of annotation maps describing the entire Gene Ontology.

26. UniProt, C. (2015) UniProt: a hub for protein information. Nucleic Acids Res, 43, D204–212.

27. van Kruijsbergen, I., Hontelez, S., Veenstra, G.J. (2015) Recruiting polycomb to chromatin. The international journal of biochemistry & cell biology, 67, 177– 187.

28. Eppig, J.T., Motenko, H., Richardson, J.E., et al. (2015) The International Mouse Strain Resource (IMSR): cataloging worldwide mouse and ES cell line resources. Mamm Genome, 26, 448–455.

29. Grozdanov, P.N., Amatullah, A., Graber, J.H., et al. (2016) TauCstF-64 Mediates Correct mRNA Polyadenylation and Splicing of Activator and Repressor Isoforms of the Cyclic AMP-Responsive Element Modulator (CREM) in Mouse Testis. Biol Reprod, 94, 34.

30. Grozdanov, P.N., Li, J., Yu, P., et al. (2018) Cstf2t Regulates expression of histones and histone-like proteins in male germ cells. Andrology.

31. Bujnicki, J.M., Feder, M., Radlinska, M., et al. (2002) Structure prediction and phylogenetic analysis of a functionally diverse family of proteins homologous to the MT-A70 subunit of the human mRNA:m(6)A methyltransferase. J Mol Evol, 55, 431–444.

